# Nascent matrix deposition supports alveolar organoid formation from aggregates in synthetic hydrogels

**DOI:** 10.1101/2024.03.19.585720

**Authors:** Madeline K. Eiken, Charlie J. Childs, Lindy K. Brastrom, Tristan Frum, Eleanor M. Plaster, Orren Shachaf, Suzanne Pfeiffer, Justin E. Levine, Konstantinos-Dionysios Alysandratos, Darrell N. Kotton, Jason R. Spence, Claudia Loebel

## Abstract

**Highlights:**

- Alveolar organoids are formed with a two-step, Matrigel-free method in a semi-synthetic hyaluronic acid (HA) hydrogel
- The two-step method offers control over alveolar size, density, and growth
- Alveolar organoids maintain their AT2 identity in HA hydrogels
- Alveolar organoids secrete nascent extracellular matrix supporting organoid growth without Matrigel

**Summary:** Human induced pluripotent stem cell (iPSC) derived alveolar organoids have emerged as a system to model the alveolar epithelium in homeostasis and disease. However, alveolar organoids are typically grown in Matrigel, a mouse-sarcoma derived basement membrane matrix that offers poor control over matrix properties, prompting the development of synthetic hydrogels as a Matrigel alternative. Here, we develop a two-step culture method that involves pre-aggregation of organoids in hydrogel-based microwells followed by embedding in a synthetic hydrogel that supports alveolar organoid growth, while also offering considerable control over organoid and hydrogel properties. We find that the aggregated organoids secrete their own nascent extracellular matrix (ECM) both in the microwells and upon embedding in the synthetic hydrogels. Thus, the synthetic gels described here allow us to de-couple exogenous and nascent ECM in order to interrogate the role of ECM in organoid formation.

## Introduction

Organoids are three-dimensional (3D) *in vitro* organ-like structures that recapitulate some elements of *in vivo* organs, including cellular diversity, spatial organization, and function (Childs et al., 2022; Clevers, 2016; Frum and Spence, 2021). Organoids representative of diverse organ systems have become powerful tools for studying a range of biological questions, which include interrogating physiologic and pathologic states to understand cell-cell and cell-matrix interactions *in vitro* (Nikolić and Rawlins, 2017; Yi et al., 2021).

Several lung organoid models, derived from primary cells or induced pluripotent stem cells (iPSCs), have been developed to study the proximal airways and distal alveoli (Barkauskas et al., 2013, 2017; Dye et al., 2015; Ekanger et al., 2022; Frum et al., 2023; Gotoh et al., 2014; Hein et al., 2022; Miller et al., 2018, 2019, 2020; Rock et al., 2009). Recently, iPSC-derived alveolar type II (AT2) organoids have emerged as a powerful tool to study alveolar homeostasis and disease such as SARS-CoV-2 (Huang et al., 2020; Hurley et al., 2020; Jacob et al., 2017; Mirabelli et al., 2021). Alveolar organoids are traditionally grown in Matrigel, a commercially available complex extracellular matrix (ECM) which supports proliferation, 3D growth, and viability (Aisenbrey and Murphy, 2020; Jacob et al., 2017). However, Matrigel, and other commercially available ECMs, are derived from non-human sources or cancer cell lines, resulting in a high biological complexity, high batch-to-batch variation, low tunability, and limited control over proteins or growth factors found within the ECM (Hughes et al., 2010; Kozlowski et al., 2021). To address these limitations, synthetic materials such as hydrogels have emerged as an alternative to commercially available complex ECMs for organoid culture (Gan et al., 2023; Hofer and Lutolf, 2021; Magno et al., 2020). Synthetic hydrogels are typically based on a polymeric backbone modified with functional moieties that provide control over the mechanical moduli (e.g., elasticity, viscosity), degradation, and the presentation of ligands to direct biological interactions (e.g., cell adhesion) (Kratochvil et al., 2019).

Several synthetic hydrogel systems have been developed for organoid culture, including hydrogel-based microwells that include an array of microcavities for aggregation and growth of single cells into organoids (Brandenberg et al., 2020; Chen et al., 2021; Choi et al., 2010; Decembrini et al., 2020; Gracz et al., 2015; Karp et al., 2007; Luan et al., 2022; Moeller et al., 2008; Wiedenmann et al., 2021). Building upon this concept, we recently developed a hydrogel-based microwell system that supports the formation and culture of alveolar organoids (Loebel et al., 2022a). Although this microwell system supports the formation of alveolar organoids with defined shape and size, it is limited for studies that aim to understand the mechanisms of organoid-hydrogel and organoid-ECM interactions.

To address this limitation, we describe a two-step culture method that leverages the advantages of microwells to control the aggregation of AT2 cells and 3D hydrogels to study organoid-hydrogel and organoid-ECM interactions. First, we demonstrated that microwell-aggregated alveolar organoids can be successfully cultured in norbornene-modified hyaluronic acid (HA) hydrogels. The HA hydrogel consists of a hyaluronic acid backbone modified with norbornenes, which enable crosslinking via a thiol-ene reaction of norbornenes with dithiols and modification with thiol moieties such as the cell-adhesive ligand RGD (Plaster et al., 2023). Using the two-step culture method, cells were first pre-aggregated in hydrogel-based microwells, then the aggregates were embedded into HA hydrogels. We demonstrated that this method offers tunability across a range of hydrogel moduli, organoid size and density and supports high organoid viability and formation efficiency, identity maintenance, and growth. Furthermore, we interrogated the mechanism of organoid formation and find that aggregates secrete their own matrix, including the basement membrane proteins laminin and collagen IV, within the microwells and after embedding in HA hydrogels, which may support organoid formation without Matrigel, highlighting the capacity of synthetic hydrogels to de-couple the role of exogenous and nascent ECM in organoid formation.

## Results

### Formation of alveolar organoids in HA hydrogels

First, we sought to test whether iPSC-derived alveolar progenitor cells form organoids within HA hydrogels fabricated with a modulus (i.e., Young’s modulus, stiffness) from 1.5-20 kPa, reflecting the Young’s moduli of alveolar regions within healthy and diseased lungs (Hinz, 2012). We compared the traditional approach of embedding single alveolar progenitor cells (400 cells/µl) into Matrigel or into HA hydrogels with initial Young’s Moduli of 1.5 kPa (soft HA hydrogels), 4 kPa (medium HA hydrogels) or 20 kPa (stiff HA hydrogels) (**Figure 1A**) (Jacob et al., 2017, 2019). The alveolar progenitor cells used here contain a tdTomato reporter targeted to the endogenous surfactant protein C (SFTPC) loci as an indicator of AT2 identity (Jacob et al., 2017, 2019). Within 10 days, single cells gave rise to alveolar organoids in Matrigel, whereas no organoids were observed within HA hydrogels (**Figure 1B, Figure S1A**). Quantification of organoid formation efficiency, defined as the number of multicellular structures present at day 14 divided by the number of aggregates at day 1, showed significant reduction for single alveolar progenitor cells in HA hydrogels (average formation efficiency: 1.3% ± 1.6%) when compared to Matrigel (formation efficiency: 4.1% ± 1.1%) (**Figure 1C**). Although single alveolar progenitor cells did not give rise to organoids in HA hydrogels, single cells remained viable throughout 14 days (average viability: 62.0% ± 19.2%), although viability was significantly lower when compared to Matrigel (97.7% ± 3.5%) (**Figure 1D**). These data indicate that single alveolar progenitor cells are not able to form organoids within HA hydrogels.

**Figure 1.**
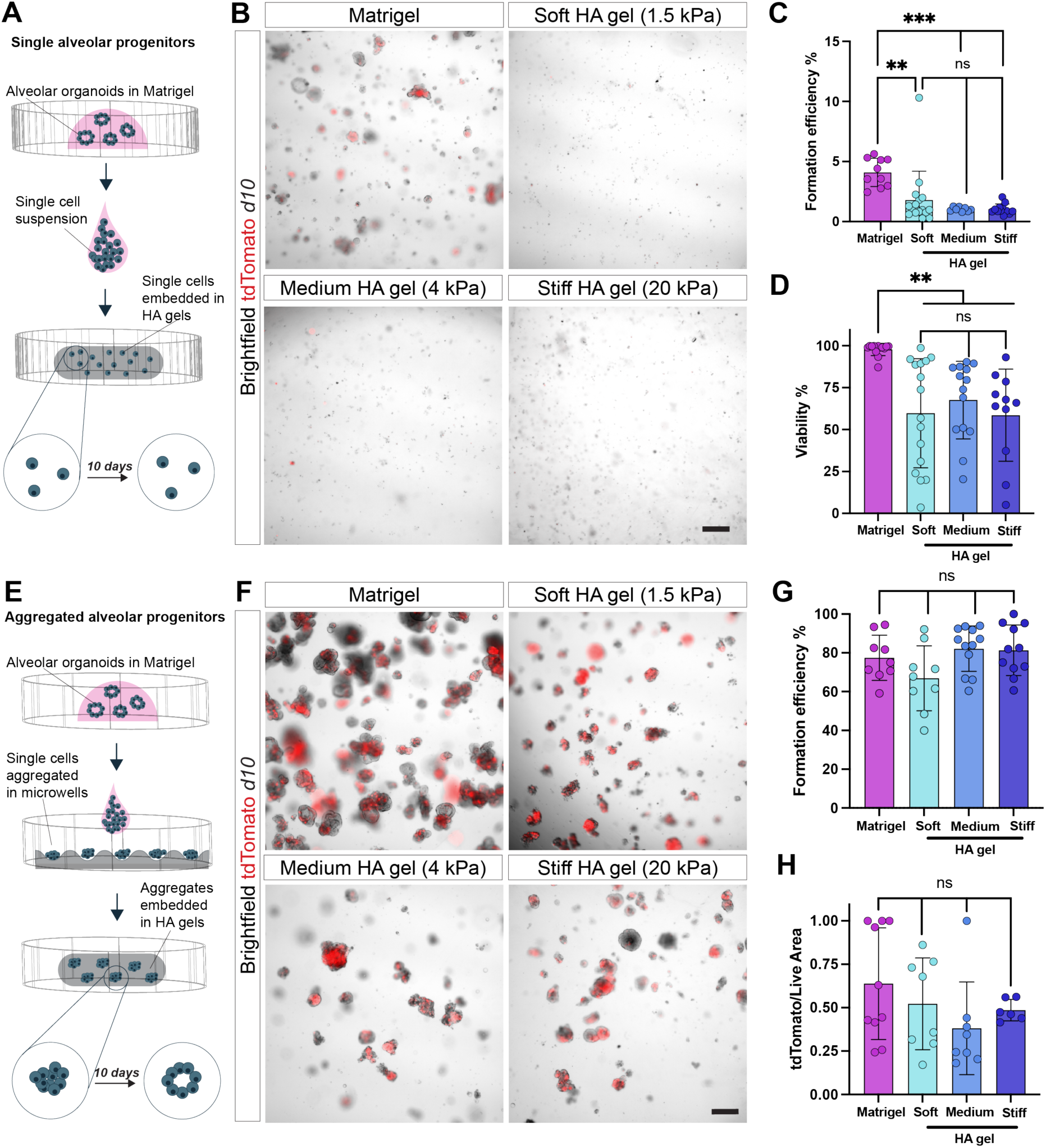
Embedding of alveolar progenitor cell aggregates enables viable alveolar organoid formation in synthetic HA hydrogels of various moduli. A. Schematic overview of experimental approach using single alveolar progenitor cells: alveolar progenitors are isolated from alveolar organoids in Matrigel and resuspended into single cells prior to embedding into HA hydrogels. B. Representative brightfield and fluorescence images showing SFTPC-tdTomato (red) expression in Matrigel and HA hydrogels with Young’s moduli of 1.5 kPa (‘soft’), 4 kPa (‘medium’) and 20 kPa (‘stiff’) at day 10 (d10). Scale bar 300 µm. C. Quantification of alveolar organoid formation efficiency of embedded single cells in Matrigel and HA hydrogels at day 14, n ≥ 8 individual hydrogels from 3 independent experiments. *** p < 0.001, ** p < 0.01, ns - no significant difference by one-way ANOVA with Tukey’s multiple comparisons test. D Viability of embedded single cells in Matrigel and HA hydrogels at day 10. n ≥ 10 regions of interest (ROI) from 3 independent experiments. ** p < 0.01, ns - no significant difference by one-way ANOVA with Tukey’s multiple comparisons test. E. Schematic overview of experimental approach using aggregated alveolar progenitor cells: alveolar progenitor cells are isolated from alveolar organoids in Matrigel, resuspended into single cells, and aggregated within HA hydrogel-based microwells for three days prior embedding into HA hydrogels. F. Representative brightfield and fluorescence images showing SFTPC-tdTomato (red) expression of aggregates in Matrigel and HA hydrogels with Young’s moduli of 1.5 kPa (‘soft’), 4 kPa (‘medium’) and 20 kPa (‘stiff’) at day 10. Scale bar 300 µm. G. Quantification of organoid formation efficiency of aggregates in Matrigel and HA hydrogels at day 14. n ≥ 8 individual hydrogels from 3 independent experiments. ns - no significant difference by one-way ANOVA with Tukey’s multiple comparisons test. H. Quantification of projected tdTomato area per live cell area (see Supplementary Figure S1C for live/dead images) of aggregates at day 10. n ≥ 6 ROI from 3 independent experiments. ns - no significant difference by ANOVA with Tukey’s multiple comparisons test.

We hypothesized that pre-aggregating alveolar progenitors within our recently developed hydrogel-based microwells will improve the formation efficiency of alveolar organoids upon embedding into HA hydrogels (Loebel et al., 2022). Dissociated alveolar progenitor cells were first seeded into microwells and allowed to aggregate for 3 days, and then embedded in Matrigel or HA hydrogels of various Young’s moduli (soft, medium, stiff) (**Figure 1E**). Within 10 days after embedding, aggregates formed into alveolar organoids in both Matrigel and HA hydrogels independent of the initial Young’s modulus (**Figure 1F**) and continued to increase in size over 14 days (**Figure S1B**). Accordingly, quantification of the organoid formation efficiency showed no significant difference between aggregates embedded in Matrigel (77.4% ± 12.4%) and HA hydrogels (average: 76.8% ± 14.9%) (**Figure 1G**). Furthermore, cell viability was maintained above 88% for aggregates embedded in Matrigel and HA hydrogels throughout the entire culture period of 14 days post-embedding (**Figure S1C**). Notably, SFTPC-tdTomato reporter expression of embedded aggregates showed no significant differences between Matrigel and HA hydrogels at day 14 (**Figure 1H**) and throughout the entire culture period (**Figure S1D**). Maintenance of SFPTC-tdTomato reporter expression was further confirmed by flow cytometry, demonstrating no significant difference in the percent of cells positive for the SFTPC-tdTomato reporter between Matrigel and HA hydrogels (**Figure S1E**). Taken together, these data show that pre-aggregation of alveolar progenitor cells within hydrogel-based microwells enhances viable alveolar organoid formation in HA hydrogels, and that HA hydrogels maintain SFTPC-tdTomato reporter expression at comparable levels to Matrigel.

### Alveolar organoids maintain AT2 identity in HA hydrogels

After showing that alveolar organoids are viable within HA hydrogels, we next aimed to characterize their identity and function. Due to minimal differences in viability between HA hydrogel stiffness, we moved forward with a medium stiffness HA gel (4 kPa), which reflects the stiffness of the healthy lung (Hinz, 2012). Given, that SFTPC-tdTomato reporter expression has limited sensitivity in detecting differences in SFTPC expression (Sun et al., 2021), we wanted to confirm that alveolar organoids maintain expression of AT2 identity markers upon embedding into HA hydrogels. Thus, we compared the transcriptional and protein expression of common AT2 cell identity markers in Matrigel and HA hydrogels. We first assessed expression of canonical AT2 genes in alveolar organoids in Matrigel and HA hydrogels using reverse transcription-quantitative polymerase chain reaction (RT-qPCR) (Frum et al., 2023; Herriges and Morrisey, 2014). No significant differences were observed for the expression of the lamellar body marker *LAMP3* and surfactant protein B (*SFTPB*) (**Figure 2A**). In contrast, expression of surfactant protein A (*SFTPA*), a common marker of AT2 maturity, and the surfactant processing protein Napsin A (*NAPSA*) were significantly lower for alveolar organoids in HA hydrogels when compared to Matrigel (**Figure 2B**). These observations suggest that AT2s are less mature in HA hydrogels. Notably, *SFTPC* expression was almost 3-fold higher in HA hydrogels when compared to Matrigel (**Figure 2C**), which we previously observed in HA microwells (Loebel et al., 2022a). Given that *SFTPC* is also expressed in less differentiated and nascent AT2 cells, these data further support the notion that HA hydrogels may prevent AT2 maturation. To assess whether HA hydrogels promote expression of non-alveolar lung markers in the organoids we also measured expression of lung development markers, including *SOX9*, *SOX2*, and *TP63*, which were overall lower for alveolar organoids in HA hydrogels (**Figure S2A**), indicating that alveolar identity is maintained in HA hydrogels. In addition, expression of the lung epithelial cell marker *NKX2.1* showed lower mRNA levels in HA hydrogels; however, similar protein expression in Matrigel and HA hydrogel argues against a functional difference between organoids grown in Matrigel and HA hydrogels (**Figure S2B**). Taken together, these transcriptional differences in the expression of AT2 markers suggest that Matrigel maintains alveolar progenitor organoids at a more mature state than HA gels.

**Figure 2.**
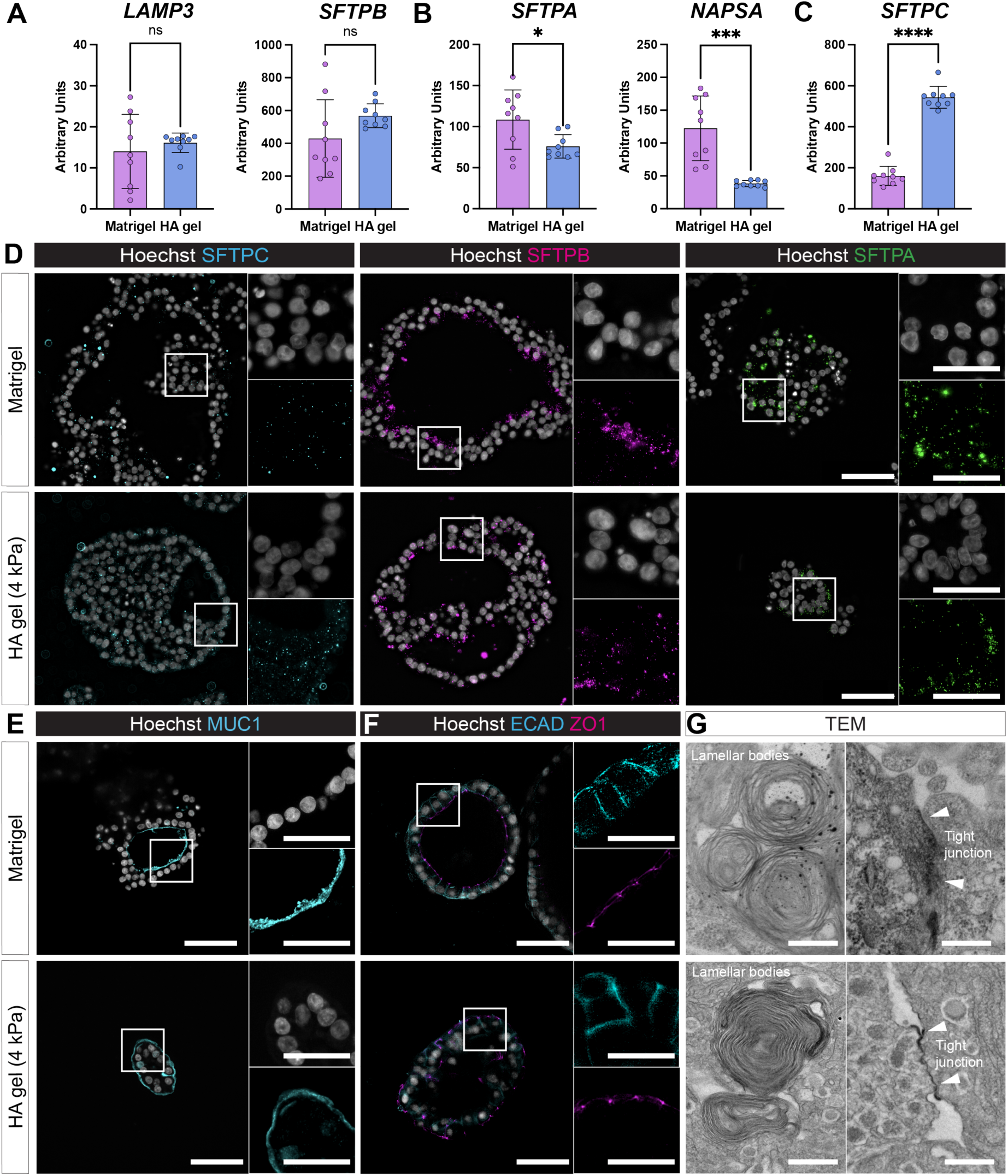
HA hydrogels maintain expression of AT2 cell identity markers of alveolar organoids. A-C. Quantification of bulk gene expression of AT2 cell identity markers, including lysosomal associated membrane protein 3 (*LAMP3*), surfactant protein B (*SFTPB*), surfactant protein A (*SFTPA*), napsin A (*NAPSA*), and surfactant protein C (*SFPTC*) for embedded alveolar progenitor cell aggregates in Matrigel and HA hydrogel (medium, 4 kPa) at day 14. n = 9 repeated qPCR measurements from 3 independent experiments. **** p<0.0001, *** p<0.001, * p<0.05, ns - no significant difference by unpaired Student’s t-test. D. Representative fluorescent images of alveolar progenitor cell aggregates embedded in Matrigel and HA hydrogel and immunostained for SFTPC, SFTPB and SFTPA and Hoechst at day 14. Scale bars 50 µm (inset 25 µm). E. Representative fluorescent images of alveolar progenitor cell aggregates in Matrigel and HA hydrogels immunostained for apical AT2 marker membrane-bound mucin (MUC1) at day 14. Scale bars 50 µm (inset 25 µm). F. Representative fluorescent images of alveolar organoids in Matrigel and HA hydrogels immunostained for epithelial cadherins (ECAD) and zonula occludens-1 (ZO1) at day 14. Scale bars 50 µm (inset 25 µm). G. Representative TEM images of organoids in Matrigel and HA hydrogels showing lamellar bodies and tight junctions (arrowheads) at day 14. Scale bars 400 nm.

Given that gene and protein expression are not always correlated (Cote et al., 2016), we next sought to confirm maintenance of AT2 identity at the protein level through immunofluorescence staining of selected AT2 proteins (Beers and Moodley, 2017; Frum et al., 2023; Jacob et al., 2017; Sucre et al., 2018; Wang et al., 2007). First, we stained embedded alveolar organoids for SFTPC, SFTPB, and SFTPA and observed the expected punctate staining patterns both in Matrigel and HA hydrogels (**Figure 2D**). In addition, AT2 cell surface markers, HTII-280 and MUC1 were examined with immunofluorescence staining and showed apical expression in alveolar organoids in both Matrigel and HA hydrogels (**Figure 2E, Figure S2C**) (Gonzalez et al., 2010; Jarrare et al., 1998), indicating cells are polarized. As expected, the apical markers were within the lumen of the organoids in Matrigel, indicating an “apical in” orientation. In contrast, apical markers were on the membrane opposite the lumen in HA hydrogels indicating an “apical out” orientation. We further interrogated organoid orientation through the presence of junctional proteins, such as epithelial cadherin (ECAD) and the tight junction protein zonula occludens-1 (ZO1). ECAD was mostly expressed at the basolateral side of the cells, whereas ZO1 was expressed as punctate spots on the apical side of cells within organoids (**Figure 2F**). Similar to the expression of AT2 cell surface markers, within Matrigel, ZO1 was mostly expressed in the lumen of the organoid, indicating an “apical-in” orientation whereas within HA hydrogels ZO1 was expressed on the outside of the organoid, indicating an “apical-out” orientation, confirming flipped epithelial cell polarity (Cereijido et al., 2008) in HA hydrogels. Finally, we confirmed AT2 identity through the presence of lamellar bodies in both Matrigel and HA hydrogels using transmission electron microscopy (TEM) (Jacob et al., 2017; Loebel et al., 2022a; Schmitz and Muller, 1991). TEM imaging also showed the expression of zipper-like lines between adjacent cells, confirming the formation of tight junctions and epithelial barrier formation (**Figure 2G**) (Buckley and Turner, 2018). Taken together, these data indicate that alveolar organoids in HA hydrogels maintain their identity as AT2 cells as shown by the expression of AT2-specific proteins, junctional complexes, and lamellar bodies.

### Initial size and density of alveolar aggregates influence organoid growth

Since we found that aggregates successfully formed organoids in HA hydrogels, we next sought to determine whether aggregate size or embedding density direct alveolar organoid growth. Our previous work demonstrated that the number of seeded cells per microwells correlates with alveolar organoid size (Loebel et al., 2022). Thus, we hypothesized that by controlling the initial size of alveolar progenitor cell aggregates we can similarly control the final alveolar organoid size after embedding in HA hydrogels (**Figure 3A**). We altered the seeding density in the microwells to include 10 cells/microwell (small), 40 cells/microwell (medium) or 120 cells/microwell (large), which all formed aggregates within 3 days. After embedding, small aggregates showed limited formation of cystic structures whereas medium and large aggregates showed an overall increase in alveolar organoid size at day 14 (**Figure 3B),** which was consistent throughout the entire culture period (**Figure S3A**). Aggregate size also impacted the growth rate of the aggregates with an average increase of 220, 345, and 700 µm^2^/organoid/day for small, medium, and larger aggregates, respectively. To assess the role of aggregate size on proliferation, we incubated cells for the first (days 0-7) or second half (days 7-14) of the culture period with 5-ethynyl 2’-deoxyuridine (EdU). Between days 0-7, the number of EdU positive nuclei increased as a function of aggregate size (**Figure 3C**). This was further confirmed by quantifying the percentage of EdU positive nuclei with a nearly 2-fold increase in EdU positive nuclei for alveolar organoids from large aggregates (34% ± 22%) when compared to those formed from small aggregates a (16% ± 18%). Interestingly, no significant difference was observed between different sized aggregates when comparing EdU expression between day 7-14 (**Figure S3B**). These data indicate that modulating the size of embedded alveolar progenitor cell aggregates provides a way to control the ultimate size of organoids, likely through differences in growth rate during the initial 7 days of culture that exaggerate initial differences in aggregate size.

**Figure 3.**
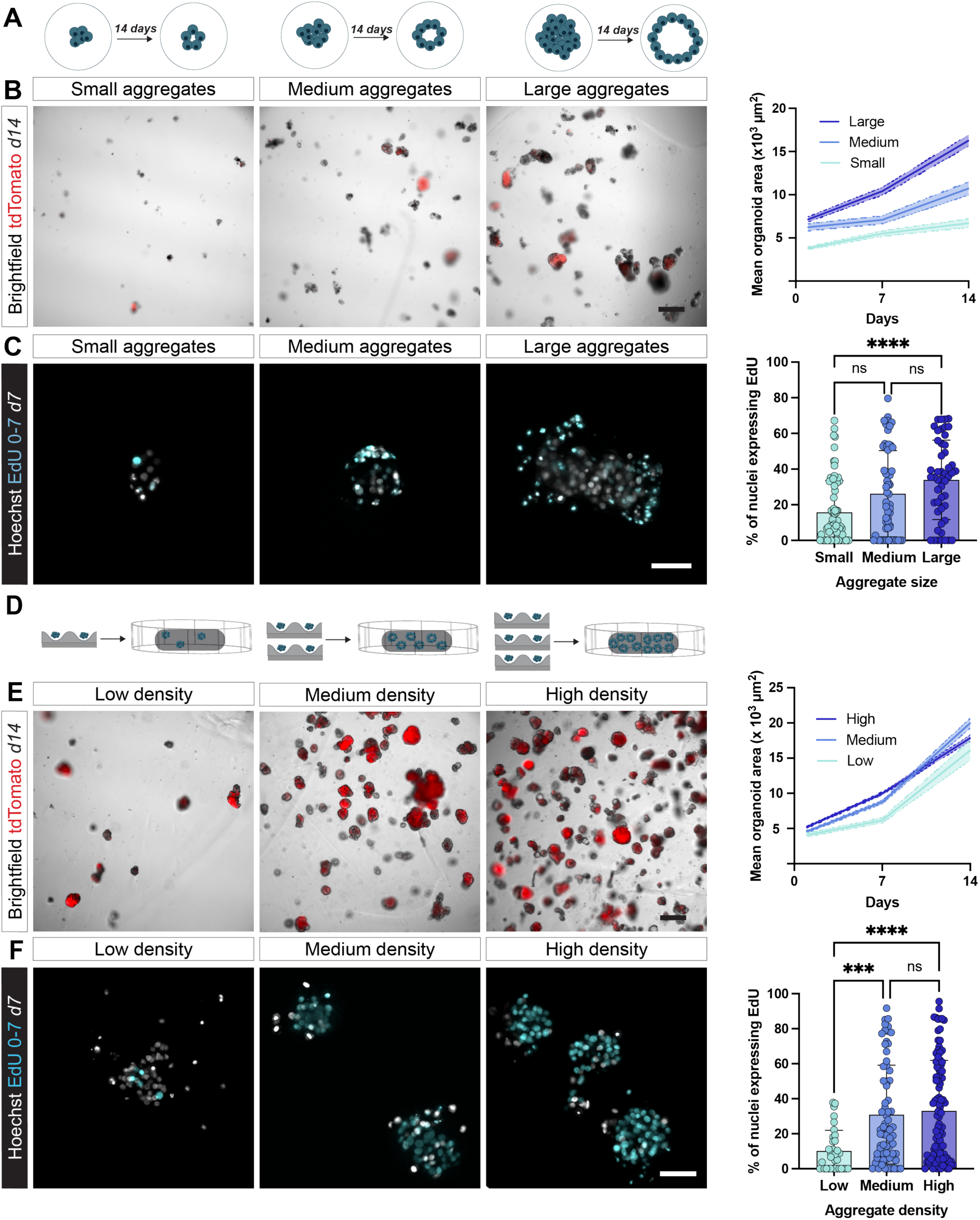
Alveolar progenitor cell aggregation enables control over alveolar organoid size and density. A. Schematic of experimental approach: the relationship between cell density in the microwell array (i.e., aggregate size) prior embedding and final alveolar organoid size. B. Representative brightfield and fluorescent images of SFTPC-tdTomato (red) expression and quantification of projected organoid size of alveolar organoids formed from small, medium, and large aggregates embedded in HA hydrogels at day 14. Scale bar 300 µm. n = 2 independent experiments. Shaded error bars represent standard error of the mean (SEM). C. Representative fluorescent images and quantification of EdU incorporation from days 0-7 of alveolar organoids formed from small, medium, and large aggregates embedded in HA hydrogels at day 7. Scale bar 50 µm. n ≥ 49 alveolar organoids from 3 independent experiments. **** p<0.0001, ns - no significant difference by one-way ANOVA with Tukey’s multiple comparisons test. D. Schematic of experimental approach: the relationship the number of embedded alveolar progenitor aggregates (medium size) and aggregate growth in HA hydrogels. E. Representative brightfield and fluorescent images of SFTPC-tdTomato (red) expression and quantification of projected organoid size of alveolar organoids formed from low, medium, and high density aggregates embedded in HA hydrogels at day 14. Scale bar 300 µm. n = 2 independent experiments.

Change in the growth rate based on aggregate size that we observed may indicate that alveolar the number of alveolar progenitor cells in proximity to each other impacts proliferation of the aggregates. To test this possibility, we varied aggregate density in the HA hydrogels by varying the number of (medium sized) alveolar progenitor cell aggregates per HA hydrogel (**Figure 3D**). Embedding of aggregates with low, medium, and high densities showed little differences in alveolar organoid structure and size at day 14; however, the density of aggregates controlled the growth kinetics (**Figure 3E**, **Figure S3C**). Alveolar organoids embedded at high and medium densities showed higher growth rates between day 0-7 of culture (700 and 800 µm^2^/day, respectively), while those embedded at low densities had slower growth rates (350 µm^2^/day). Here as well, incorporation of EdU confirmed these findings with an almost 300% increase in EdU expression for alveolar organoids formed from aggregates embedded at medium and high densities (31% ± 28% and 33% ± 29%, respectively) when compared to aggregates embedded at low densities (10% ± 12%) (**Figure 3F**). No significant differences in EdU incorporation were observed between day 7-14 (**Figure S3D**). These data indicate that alveolar progenitor cells support the growth of each other, at both the cell-cell scale within organoids, based on the aggregate size experiment, and at longer scales between organoids, based on the aggregate density experiment. In addition, it shows that our two-step culture method enables control over the size and density at which alveolar aggregates are embedded and thus, provides a useful tool to study the role of cell signaling within and across organoids. Shaded error bars represent SEM. **F**. Representative fluorescent images and quantification of EdU incorporation from days 0-7 of alveolar organoids formed from low, medium, and high density aggregates embedded in HA hydrogels at day 7. Scale bar 50 µm. n ≥ 34 alveolar organoids from 3 independent experiments. **** p<0.0001, *** p<0.001, ns - no significant difference by one-way ANOVA with Tukey’s multiple comparisons test.

### Aggregates secrete nascent ECM in microwells and upon embedding in HA hydrogels

After having shown that cellular interactions are important for organoid growth in HA hydrogels, we next sought to investigate the interactions between cells and their surrounding ECM. Building upon previous work showing ECM deposition by epithelial organoids (Blatchley et al., 2022), we hypothesized that aggregates start to secrete components of the ECM within the microwells (**Figure 4A**). To interrogate nascent ECM secretion, we leveraged a glycan labelling approach that is based on the incorporation of a galactosamine analog (Ac_4_GalNAz) into nascent glycans. Upon secretion, Ac_4_GalNAz-containing glycans are modified with a fluorophore using click-chemistry (Dube et al., 2006; Laughlin et al., 2008; Zhang and Zhang, 2013). In addition to nascent glycans, we stained for the basement membrane proteins, laminin, and collagen IV (**Figure 4B**), which are known to be secreted by lung epithelial cells (Nguyen et al., 2002; Pierce et al., 1998; Rosmark et al., 2023). Nascent glycan expression was largely confined to the cell membranes, likely labeling o-linked mucins or glycosphingolipids in the glycocalyx on the cell surface (Aich and Yarema, 2008). In contrast, aggregates were surrounded by laminin, whereas collagen IV was deposited towards the center of the aggregate as punctate spots. Given that collagen IV but not laminin contains galactosamine in the globular domain (Weber et al., 1984), collagen IV staining showed some overlap with the nascent glycan staining (**Figure S4A**). Importantly, no exogenous laminin or collagen IV were added to the cultures. Thus, these findings show that alveolar progenitor cells secrete nascent ECM that forms a shell around the aggregates in the microwells. Based on this observation, we next hypothesized that organoid continue to secrete nascent ECM upon embedding in the HA hydrogels (**Figure 4C**). To demonstrate this, we continued nascent ECM labeling after embedding aggregates into HA hydrogels and throughout the 14 days of culture (**Figure S4B**). Over the first few days, nascent glycans remained largely confined to the cell membrane but showed a significant increase in overall glycan thickness on day 14 (**Figure S4C, Figure 4D**). These observations suggest that alveolar organoids continued to secrete nascent ECM upon embedding within HA hydrogels. Previous studies have shown that nascent ECM deposition within synthetic hydrogels plays a synergistic role in directing cell function (Ferreira et al., 2018; Loebel et al., 2019). Thus, we next wanted to assess whether nascent glycans influence the direct interactions of alveolar organoids with the HA hydrogel. To quantify the organoid-hydrogel distance, we co-embedded fluorescent microbeads (0.5 µm) to visualize the hydrogel (Loebel et al., 2020) (**Figure S4D**). Similarly to the nascent glycan thickness, the organoid-hydrogel distance, as measured by the distance between the cell membrane and the microbeads, increased over the 14 days of culture **(Figure 4E)**. In addition, we did not find any fluorescent microbeads within the nascent glycans (**Figure S4E**), suggesting that the nascent ECM is forming an interface between organoids and the HA hydrogels. We also stained for laminin and collagen IV that showed constant (laminin) or decreasing (collagen IV) secreted by day 14 when normalized to organoid size, which is likely due to organoid growth (**Figure 4F, Figure S4G**). Taken together, these data confirm our previous findings that nascent ECM supports cell function by forming an interface between cells and the synthetic hydrogel.

**Figure 4.**
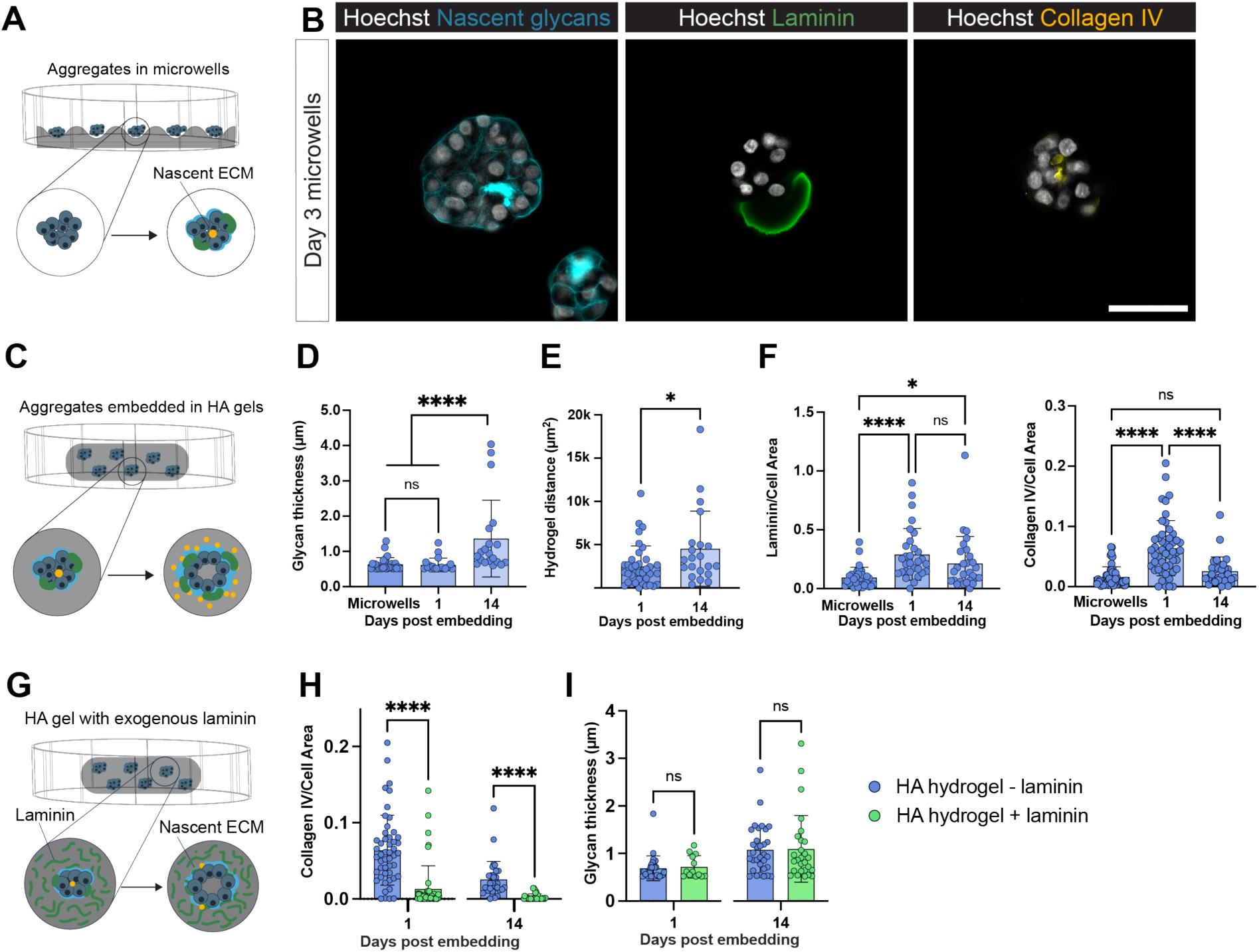
Nascent ECM of alveolar progenitor cell aggregates supports alveolar organoids A. Schematic of experimental approach: ECM deposition by aggregates in microwells was visualized. B. Representative fluorescent images of ECM secreted by the aggregates in the microwells (day 3). Scale bar 50 µm. C. Schematic of experimental approach: ECM deposition by aggregates/organoids upon embedding into HA hydrogels was visualized and quantified. D. Quantification of projected nascent glycan thickness beyond the cell membrane across the culture period. n ≥ 20 alveolar organoids from 3 independent experiments. **** p<0.0001, ns - no significant difference by one-way ANOVA with Tukey’s multiple comparisons test. E. Quantification of the organoid-hydrogel distance (i.e. the area between the aggregate/organoid and hydrogel after embedding aggregates in HA hydrogels) across the culture period. n ≥ 20 alveolar organoids from 3 independent experiments. * p<0.05, ns - no significant difference by unpaired Student’s t-test. F. Quantification of projected laminin and collagen IV per aggregate/organoid area. n ≥ 28 aggregates/organoids from 3 independent experiments. **** p<0.0001, * p<0.05, ns - no significant difference by one-way ANOVA with Tukey’s multiple comparisons test. G. Schematic of experimental approach: laminin was co-embedded in the HA hydrogels with aggregates, and subsequent collagen IV and nascent glycan secretion by the aggregates was assessed. H. Quantification of collagen IV per cell area in HA hydrogels without laminin (blue bars) and with added laminin (green bars) at day 1 and day 14. n ≥ 31 ROI across 3 independent experiments. *** p<0.001, **** p<0.0001 by unpaired Student’s t-test. Scale bars 50 µm. I. Quantification of projected nascent glycan thickness in HA hydrogels with no additional proteins (blue bars) and HA hydrogels with laminin (green bars). N ≥ 13 ROI across 2 independent experiments. ns - no significant difference by unpaired Student’s t-test.

Given our findings that organoid aggregates produce nascent ECM in the absence of exogenous ECM, coupled with prior studies showing that addition of exogenous ECM proteins are necessary for organoid formation from single cells in synthetic hydrogels (Broguiere et al., 2018; Gjorevski et al., 2016; Loebel et al., 2022), we wanted to determine if the lack of exogenous ECM in our system promotes nascent ECM deposition (**Figure 4G**). To test this, we compared nascent glycan and laminin deposition of alveolar organoids within HA hydrogels without or with laminin/entactin (2 mg/ml) mixed into the hydrogel. We observed a significant decrease in the expression of collagen IV in alveolar organoids cultured within HA gels with added laminin (HA hydrogel + laminin) when compared to HA hydrogels without additional laminin (**Figure 4H, Figure S4G**). Although not significant, collagen IV mRNA was also reduced for alveolar organoids in HA/laminin hydrogels (**Figure S4H**). There was no significant difference in nascent glycan thickness in the HA hydrogels with or without laminin (**Figure 4I, Figure S4I**). These data show that the presence of exogenous ECM basement membrane proteins reduces the need for alveolar progenitors to deposit their own ECM in HA hydrogels. In summary, microwells support the formation of organoid aggregates, which, in the absence of exogenous ECM, produce nascent ECM which may support organoid formation and growth within synthetic hydrogels.

## Discussion

Here, we described a two-step culture method for the formation of alveolar organoids. Our method includes the pre-aggregation of dissociated cells in microwells, followed by embedding in an HA hydrogel, which supports high viability and growth of AT2 organoids as well as maintenance. Importantly, the microwells in the pre-aggregation step provide precise control and tunability over aggregate size when compared to the traditional method of forming organoids in Matrigel, which gives rise to highly heterogeneous population of organoids. In addition, the ability to control the seeding density in HA hydrogels provides means to control cell-cell and organoid-organoid signaling, which are critical factors in directing organoid growth. Our findings further show that single cells cannot make viable organoids when directly embedded in hydrogels, but that pre-aggregated organoids may have a growth advantage in HA hydrogels because they deposit their own ECM, supporting their formation and growth upon embedding in HA hydrogels. This two-step method also enables culture of alveolar organoids in HA hydrogels of various moduli, which may provide a useful tool to study how matrix stiffness and organoid-matrix interactions direct alveolar progenitor cell function (Ahmed et al., 2023; Liu and Tschumperlin, 2011), currently not possible in traditional Matrigel cultures.

Interestingly, our study showed that pre-aggregated organoids grow with an ‘apical-out’ orientation in HA hydrogels in contrast to the typical ‘apical-in’ orientation within Matrigel as shown by HTII280, MUC1, and ZO1 staining (**Figure 2E,F, Figure S2C**). Such flipped orientations have been observed in other organoids in the absence of Matrigel (Capeling et al., 2019, 2022; Co et al., 2019, 2021; Fiorotto et al., 2023; Kakni et al., 2022), and is thought to be due to the basement membrane proteins present in Matrigel that direct polarity through integrins. Given that cell polarity is largely directed by integrin binding to the ECM (Streuli, 2009), the presence of laminin and collagen IV in Matrigel likely guides cellular orientation and the ‘apical-in’ polarization with the apical side of the organoids facing away from the Matrigel. It is important to note that we observed this ‘apical-out’ orientation in the HA hydrogels despite the presence of Arg-Gly-Asp (RGD), a fibronectin-derived integrin binding domain (Brandley and Schnaar, 1988), which may be explained by the fact that fibronectin is not part of the basement membrane. While laminin is well-known to direct epithelial cell polarity (Matlin et al., 2017), deposited laminin within microwells that was embedded with the aggregates had no effect on the ‘apical-out’ orientation upon HA hydrogel embedding (**Figure 4B**). Indeed, this has been described in other studies showing that laminin alone may not be sufficient in re-orienting organoid polarity (Martins-Costa et al., 2023). On the other hand, we previously observed an ‘apical-out’ orientation of organoids after long-term culture in microwells (Loebel et al., 2022a). Thus, it is possible that the collagen IV in the center of the aggregates (**Figure 4B**) guides their orientation as they form into organoids within the HA hydrogel. It has also been suggested that once polarity is established, cells may not be able to re-orient themselves in all organoid models (Wijesekara et al., 2019). A potential re-orientation in response to nascent ECM deposition is also limited by the capacity of cells to remodel the relatively stiff HA hydrogels that are not easily degradable by embedded organoids. In summary, our two-step culture method provides a useful platform to further study the role of the nascent ECM in guiding organoid polarity.

Another critical component of this study is the introduction of click chemistry to label nascent glycans that are produced by aggregates and organoids. Nascent glycan labeling has been used extensively in glycobiology (Dube et al., 2006; Saxon and Bertozzi, 2000; Zhang and Zhang, 2013) and the galactosamine analog has specifically been used to label nascent glycans in developing zebrafish (Laughlin et al., 2008). Although, click chemistry approaches have been used for directing assembly of cell spheroids (O’Brien et al., 2015), and nascent ECM staining has been used to visualize the cell-hydrogel interface of single cells embedded in synthetic hydrogels (Günay et al., 2023; Loebel et al., 2019), it has not been used to study the cell-hydrogel interface of complex multicellular structures such as organoids. It is well-established that cells remodel their environment both *in vivo* and *in vitro*. Multiple cell types have been shown to deposit ECM in synthetic hydrogels, including stromal cells such as mesenchymal stromal cells (MSCs) (Ferreira et al., 2018; Hezaveh et al., 2018; Loebel et al., 2019), fibroblasts (Ivarsson et al., 1997; Mochitate et al., 1991; Zhou et al., 2014), endothelial cell cocultures with fibroblasts (Friend et al., 2023; Juliar et al., 2020; Moon et al., 2010; Peters et al., 2016), chondrocytes (Bryant and Anseth, 2002; Kisiday et al., 2002; Richardson et al., 2019), and oocytes (Tomaszewski et al., 2021). In addition, ECM deposition by cells in hydrogels have important implications for the way that cells can sense and interact with their surrounding environment (Loebel et al., 2019). Although epithelial cells secrete ECM (Qureshi et al., 2017; Rosmark et al., 2023; Van Der Velden et al., 2018), the deposition of matrix has been minimally explored in the context of epithelial cells in hydrogels or epithelial organoids in hydrogels, though recent papers have explored the role of ECM deposition in organoids containing stromal cells (Below et al., 2022; Güney et al., 2021) and primary tissue-derived organoids (Fiorotto et al., 2023). One important aspect that requires additional investigations is the modification of synthetic hydrogels with cell-adhesive peptides. For example, some required the use of RGD in synthetic hydrogels to support organoid formation and growth (Cruz-Acuña et al., 2017; Gjorevski et al., 2016) while others showed successful organoid formation in alginate hydrogels without additional cell adhesive ligands (Capeling et al., 2019), perhaps due to the presence of mesenchymal cells that produced an adequate amount of ECM. Finally, our study showed that pre-aggregation of epithelial cells allows them to deposit a supportive ECM layer that supports the formation and growth of organoids within 3D synthetic hydrogel. The fact that organoids synthesize and deposit little endogenous collagen IV in laminin-containing HA hydrogels supports the notion that epithelial organoids sense and respond to environmental cues to help establish their own niche. Overall, this study uses a two-step culture method to support Matrigel-free alveolar organoid culture, highlighting the importance of both cell-cell and cell-ECM interactions in organoid formation and growth, and value of synthetic hydrogels in enabling the study of organoid hydrogel-interactions.

## Experimental Procedures

### Resource availability

*Corresponding author* Claudia Loebel

#### Materials availability

Materials are described in the main text or supplemental information. Requests for additional data or materials can be made to the corresponding author. Data and materials will be shared based on material transfer agreements.

### iAT2 generation and maintenance

iAT2 cells were differentiated from iPSCs as previously described and procured as mature iAT2s from the Center for Regenerative Medicine’s repository at Boston University (Jacob et al., 2017). Cells were received at day 120 and used in experiments up to day 300. Cells were cultured in Iscove’s Modified Dulbecco’s Medium (IMDM)/Ham’s F12 media supplemented with CHIR99021 (3 µg), keratinocyte growth factor (KGF) (10 ng/ml), dexamethasone (50 µM), 3-isobutyl-1-methylxanthine (IBMX) (0.1 mM), 8-bromo-cAMP (43 µg/ml), and primocin (called CK+DCI medium). For the initial 48 hours in culture, the media contained Y-27632 (Y27) (10 µM, Tocris Catalog #1254); after this, cells were further cultured in CK+DCI. Alveolar organoids were then passaged every 10 days by mechanically dissociating Matrigel-embedded alveolar organoids (Corning, Catalog #354234) with a pipette tip, followed by treatment with a 0.05% Trypsin/EDTA solution (Gibco Catalog #225200-054), and embedding into 50 µL Matrigel droplets at a density of 400 cells/µl. Droplets were then incubated at 37 °C and 5% CO_2_ for 20 minutes to enable crosslinking followed by adding the media. For cryopreservation, cells were dissociated using the protocol above, then suspended in a cryopreservation solution of 10% DMSO (Sigma Catalog #D2650-100ML) and 90% FBS (Corning Catalog #MT35015CV). Cells were thawed by seeding in Matrigel at high density (500,000 cells per Matrigel droplet) in the same media conditions used after passaging. Cells were confirmed to be sterile and mycoplasma-free prior to beginning the study, and periodically sorted to purify for the SFTPC-tdTomato positive population.

### Hydrogel preparation and cell encapsulation

#### Hydrogel synthesis

Norbornene-modified HA was prepared as previously described (Loebel et al., 2019) Norbornene modification degree was confirmed by ^1^H NMR to be 26% (**Figure S5A, B**). Di-thiolated peptides that are susceptible to metalloproteinase (MMP) cleavage (GCNS*VPMS*MRGGSNCG) and thiolated cell-adhesive peptides (GCGYG*RGD*SPG) were obtained from Genscript. HA hydrogels were fabricated by mixing 4 wt% norbornene-modified HA with 0.05 wt% photo-initiator lithium phenyl-2,4,6-trimethylbenzoylphosphinate (LAP, Colorado Photopolymer Solutions) and crosslinked for 3 minutes with either ultraviolet light (UV, 320-390 nm, Omnicure S1500, Exfo) for microwells, or visible light (400-500 nm, Omnicure S1500, Exfo) for gels with embedded cells or aggregates. HA hydrogel mechanics (elastic modulus) was assessed using shear rheometry (Plaster 2023) (**Figure S5C, D**).

#### Microwell generation

Microwell hydrogels were generated as previously described (Loebel 2022). Briefly, microwells were molded from cell culture microwell plates (Nacalai Catalog #4000-9VP, diameter: 400-500 µm, depth: 100-200 µm). First, negative molds were generated from poly(dimethylsiloxane) (PDMS, Sylgard 184, procured from Ellsworth Adhesives with a 9:1 base:crosslinker ratio), mixed with Hexanes (Fisher Scientific H292-1) in a 30% vol/vol ratio, incubated in a fume hood for 30 min, followed by crosslinking for 2 hours at 80°C. Microwell hydrogels (Young’s Modulus of 30 kPa) were then fabricated atop PDMS molds using methacrylated coverslip as previously described (Loebel et al., 2022). Following sterilization of microwell hydrogels within a germicidal UV light box, microwells were washed with PBS, then single cell suspensions of iAT2 cells were seeded at varying densities and cultured for three days in CK+DCI media supplemented with 10 µM Y27.

#### Aggregate encapsulation

After three days of microwell hydrogel culture, iAT2 aggregates were lifted off the microwells using PBS and centrifuged at 200 rcf, iAT2 aggregate pellets were then mixed with hydrogel precursor solutions (4 wt% norbornene-modified HA), 1 mM RGD, MMP crosslinkers (0.4-5.0mM), pipetted between two coverglasses with a 300 µm spacer, and crosslinked with visible light (5 min, 13 mW/cm^2^) for 5 minutes. For studies investigating the role of exogenous laminin, 2 mg/ml laminin-entactin complex (Corning Catalog #354259) was additionally mixed with the hydrogel precursor prior crosslinking. After crosslinking, HA hydrogels were cut into 4 x 4 mm pieces and cultured in CK+DCI medium supplemented with 10 µg/ml Y27 for the first 48 hours, followed by CK+DCI media for up to 14 days with media changes every 2-3 days.

### Cell analyses

#### Cell viability

Cell viability was assessed by incubating Matrigel and HA hydrogel embedded aggregates with Hoechst (1:1000, Thermo Fisher Scientific Catalog #33342), Calcein AM (1:2000, Invitrogen Catalog #C3099), and NucRed Dead 647 (2 drops per 1 ml, ThermoFisher Catalog #R37113) for 30 min at 37°C, followed by imaging with a Leica THUNDER microscope. Quantification of cell viability (live cell area (Calcein AM positive) per total cell area (live cells + dead cells (NucRed Dead 647 positive)) was performed using ImageJ (U. S. National Institutes of Health, Bethesda, Maryland, USA, https://imagej.nih.gov/ij/).

#### Flow cytometry

Alveolar organoids were removed from Matrigel by mechanical dissociation with a cut pipette tip or retrieved from HA hydrogels with 2% hyaluronidase (Sigma Catalog #37326-33-3) for 30 minutes at 37°C followed by mechanical dissociation with a pipette tip. Aggregates were washed twice with 1 ml PBS and disassociated into single cells (10 minutes 37°C in 0.05% Trypsin/EDTA). The single cell suspension was washed with 1 ml of FACS buffer (PBS, 2% BSA and 10 µM Y-27), followed by filtering through a 70 µM tube-top filter. A 1:2000 dilution of DAPI was added to cells in FACs buffer. FACS was performed on a Sony MA900.

#### RNA extraction, reverse transcription, and RT-qPCR

Alveolar organoids were removed from Matrigel and HA hydrogels as described above, pelleted, and flash frozen by submerging in liquid nitrogen. RNA was isolated from the frozen pellets following the instructions provided by the manufacturer (MagMax-96 Total RNA Isolation kit, Thermo Fisher Catalog #AM1830), and concentrations measured with a Nanodrop 2000. cDNA synthesis was performed using 100 ng RNA from each sample and using the SuperScript VILO cDNA Kit (Thermo Fisher Scientific, Catalog #11754250). RT-qPCR was performed on a Step One Plus Real-Time PCR System (Thermo Fisher Scientific, Catalog #43765592R) using QuantiTect SYBR Green PCR Kit (QIAGEN, Catalog #204145). Expression of genes in the measurement of arbitrary units was calculated relative to RN18S using the following equation: 2^RN18S(CT)−GENE(CT)^ × 1,000. See **Supplementary Table 1** for primers.

#### Immunofluorescence identity

HA hydrogel embedded alveolar organoids were fixated in 4% paraformaldehyde (ThermoFisher Scientific Catalog #J61899) for 30 minutes at 4°C, followed by incubation in 2 wt% hyaluronidase as described above. Matrigel embedded alveolar organoids were mechanically dissociated from the Matrigel. Organoids were subsequently embedded in Histogel (Fisher Scientific Catalog #HG-4000-012) and dehydrated via a graded alcohol series (one hour per step): 25% methanol (Sigma Aldrich Catalog #179337), 50% methanol, 75% methanol, and 100% methanol. After dehydration, alveolar organoids were soaked in 100% ethanol (Decon Labs Catalog #2701), 70% ethanol, and then infiltrated with paraffin using an automated tissue processor (Leica ASP300) overnight. 5 μm sections were then cut from the paraffin blocks and placed onto charged glass slides and stored at room temperature (RT) with a silica desiccator packet. Next, slides were baked for an hour in a 60°C dry oven, rehydrated two times for 5 minutes using Histo-Clear II (National Diagnostics, Catalog #HS-202), followed by two times rinsing in PBS for 2 minutes and a series of solutions (5 minutes per step): 100% ethanol, 95% ethanol, 70% ethanol, 30% ethanol, and double-distilled water (ddH_2_O). Antigen retrieval was performed using 1× sodium citrate buffer (100 mM trisodium citrate MilliporeSigma Catalog # S1804) and 0.5% Tween 20 and two rinses in ddH2O. Humidity chambers at RT were used for blocking and incubating the solution (5% normal donkey serum (MilliporeSigma Catalog #D9663) with PBS and 0.1% Tween 20. The slides were then incubated in primary antibodies diluted in the blocking solution overnight at 4°C in the humidity chamber. Subsequently, the slides were washed in 1× PBS three times for 5 minutes and incubated with secondary antibodies and Hoechst (1:1000) diluted in a blocking solution for one hour at RT in the humidity chamber. After three washes in 1× PBS for 5 minutes, slides were mounted with ProLong Gold (Thermo Fisher Scientific Catalog #P369300) and imaged within a week on a Leica THUNDER microscope. See **Supplementary Table 2** for antibodies and dilutions.

#### EdU staining

Aggregates were embedded in HA hydrogels as described above and cultured in CK+DCI media containing 5 µM EdU for the described culture periods. Upon fixation, incorporated EdU was labeled using the Click-&-Go Plus EdU Cell Proliferation Kit for Imaging (Click Chemistry tools Catalog #1314) and imaged with a Leica THUNDER microscope. EdU incorporation was quantified by the number of EdU positive nuclei per total number of nuclei.

#### Metabolic labeling

For nascent protein analysis, aggregates were embedded as described above and cultured in CK+DCI media with 50 µM azide-modified galactosamine (Click Chemistry Tools Catalog #2281). Click-labeling of nascent proteins was performed with 8.3 µM dibenzocyclooctyne (DBCO)−488 (Click Chemistry Tools Catalog #1278-5) in PBS for 30 minutes at 4°C, followed by fixation with 4% paraformaldehyde for 30 min at 4°C. In addition, cells were stained with Hoechst (1:1000) and CellMask deep red (1:1000) (Invitrogen Catalog #C10046) or phalloidin 647 (1:400) (Invitrogen Catalog #2326023) for 60 min at room temperature. Images were obtained with a Leica THUNDER microscope. Quantification of nascent protein thickness was performed using the ImageJ plugin BoneJ plugin as previously described (Loebel et al., 2022).

#### Immunofluorescence staining of cell-cell contacts and ECM proteins

Upon fixation, alveolar organoids were retrieved from HA hydrogels and embedded into HistoGel as described above, followed by permeabilization in 0.1% Triton-X (Sigma-Aldrich Catalog #T8787) for 40 min at 4°C and blocking with 2 wt% bovine serum albumin (BSA) for 1 hour at RT. Samples were incubated in primary antibodies for 48 hours in 2 wt% BSA at 4 °C, followed by 1 h incubation at RT in secondary antibodies and Hoechst (1:1000). Staining for ECM proteins was performed without permeabilization. All images were obtained with a Leica THUNDER microscope and quantified using ImageJ. Nascent protein area was reported as specific protein area per organoid cell membrane area.

#### Transmission Electron Microscopy (TEM)

TEM preparation and imaging of HA-embedded alveolar organoids hydrogels was performed by the University of Michigan BRCF Microscopy and Image Analysis Laboratory. Samples were first fixated in 3% glutaraldehyde + 3% paraformaldehyde in 0.1 M cacodylate buffer (CB), pH 7.2. Samples were washed 3 times for 15 minutes in 0.1 M CB, then processed for 1 hour on ice in a post-fixation solution of 1.5% K_4_Fe(CN)_6_ + 2% OsO_4_ in 0.1 M CB. Samples were then washed 3 times in 0.1 M CB, and 3 times in 0.1 M Na_2_ + acetate buffer, pH 5.2, followed by en bloc staining for 1 hour in 2% uranyl acetate + 0.1 M Na_2_ + acetate buffer, pH 5.2. Samples were then processed overnight in an automated tissue processor, including dehydration from H_2_O through 30%, 50%, 70%, 80%, 90%, 95%, 100% EtOH, followed by 100% acetone. Samples were infiltrated with Spurr’s resin at a ratio of acetone/Spurr’s resin of 2:1 for 1 hour, 1:1 for 2 hours, and 1:2 for 16 hours, then absolute Spurr’s resin for 24 hours. After embedding and polymerization, samples were sectioned on an ultramicrotome (Leica). Imaging was performed on a JEOL JEM 1400 Plus TEM.

### Statistical Analysis and Reproducibility

GraphPad Prism 9 software was used for statistical analyses. Statistical comparisons between two experimental groups were performed using unpaired Student’s *t*-tests and comparisons among more groups were performed using one-way analysis of variance (ANOVA) with Tukey’s multiple comparisons test. All experiments were repeated as described in the text.

## Acknowledgements

CL is supported by the National Heart, Lung, and Blood Institute (NHLBI; R00-HL151670) and by the American Lung Association (IA-939940). JRS is supported by the Cystic Fibrosis Foundation Epithelial Stem Cell Consortium, by the Chan Zuckerberg Initiative, an advised fund of the Silicon Valley Community Foundation (CZF2019-002440), and by the National Heart, Lung, and Blood Institute (NHLBI; R01-HL166139). MKE is supported by the National Science Foundation Graduate Research Fellowship (DGE-2241144). Human iPSC-derived AT2 cells were received from the Center of Regenerative Medicine of Boston University and Boston Medical Center, NIH/NHLBI grants N01: 75N92020C00005 and R01HL095993 to DNK. Any opinion, findings, and conclusions or recommendations expressed in this manuscript are those of the authors. The authors are grateful for support from the University of Michigan BRCF Microscopy and Image Analysis Laboratory, the University of Michigan’s BRCF Flow Cytometry Core, and Dr. Minli Xing from the University of Michigan BioNMR (U-M BioNMR) Core Facility (supported by the U-M College of Literature, Sciences, and Arts, the Life Sciences Institute, the College of Pharmacy, and the U-M Biosciences Initiative).

## Author contributions

MKE, CL, and JRS designed experiments. TF provided critical expertise and experimental input. MKE, CJC, LKB, EP, SP performed experiments and collected data. MKE, CJC, OS, and JEL analyzed data. DNK and KDA provided critical materials (alveolar organoids). MKE, CL, and JRS interpreted data and wrote and prepared the manuscript. All authors contributed to manuscript review and editing.

## Declaration of Interests

DNK holds intellectual property relating to alveolar organoids. JRS and TF hold intellectual property related to lung organoid technologies. The remaining authors declare no competing interests.

