## Supplement for "Nascent matrix deposition supports alveolar organoid formation from aggregates in synthetic hydrogels"

[1] Department of Biomedical Engineering, University of Michigan, Ann Arbor, USA
[2] Department of Cell and Developmental Biology, University of Michigan Medical School, Ann Arbor, MI, USA
[3] Department of Computational Medicine and Bioinformatics, University of Michigan Medical School, Ann Arbor, MI, USA
[4] Department of Internal Medicine, University of Michigan Medical School, Ann Arbor, MI, USA
[5] Department of Biomedical Engineering, University of Texas, Austin, TX, USA
[6] Center for Regenerative Medicine, Boston University and Boston Medical Center, Boston, MA 02118, USA
[7] The Pulmonary Center and Department of Medicine, Boston University Chobanian & Avedisian School of Medicine, Boston, MA 02118, USA
[8] Program in Cell and Molecular Biology, University of Michigan Medical School, Ann Arbor, MI, USA
[9] Department of Materials Science and Engineering, University of Michigan, Ann Arbor, MI, USA

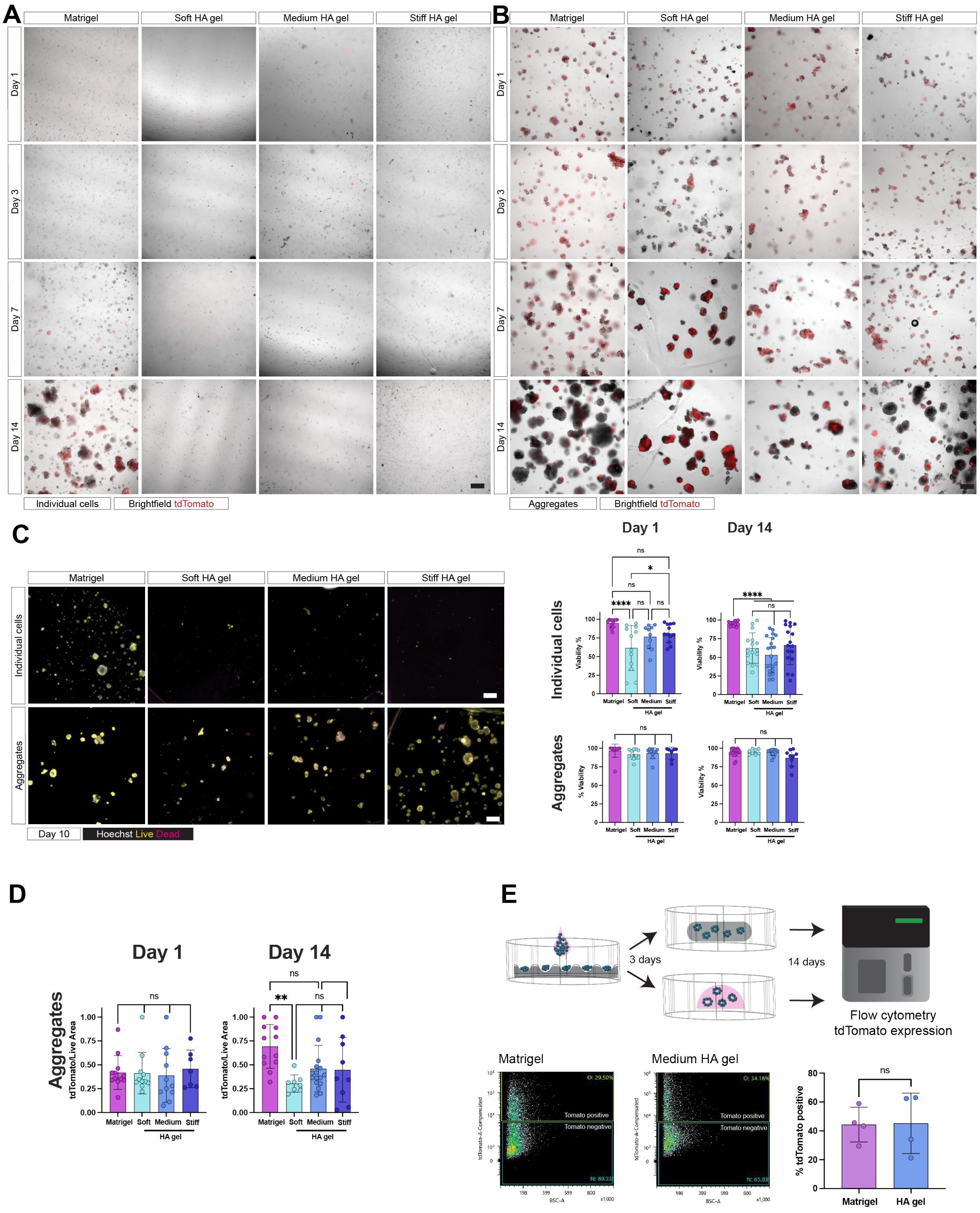


**Figure S1. Aggregates form in HA hydrogels and maintain reporter expression**. **A**. Representative images of individual cells in HA hydrogels for up to 14 days. Scale bar 300 µm. **B**. Representative images of aggregates in HA hydrogels for up to 14 days. Scale bar 300 µm. **C**. Representative images of live (yellow) and dead (magenta) cells for viability quantification of individual cells and aggregates at day 10 in culture. Scale bar 50 µm. n ≥ 6 regions of interest from 3 independent experiments. **** p<0.0001, * p<0.05, ns – no significant difference by one-way ANOVA with Tukey's multiple comparisons test. **D**. Quantification of SFTPC-tdTomato area per live area of aggregates. n ≥ 6 ROI from 3 independent experiments. ** p<0.01, ns – no significant difference by one-way ANOVA with Tukey's multiple comparisons test. **E**. Flow cytometry experiment to confirm SFTPC-tdTomato reporter expression in Matrigel and HA hydrogels. Experimental schematic: cells are aggregated and embedded either in Matrigel or HA hydrogels, followed by quantification of td-Tomato expression by flow cytometry. Representative flow plots of cells embedded in Matrigel and HA hydrogels, showing tomato positive cells (box O) and tomato negative cells (box N). tdTomato expression is based on 4 independent experiments in Matrigel and HA hydrogels. ns - no significant difference by unpaired Student's *t*-test.

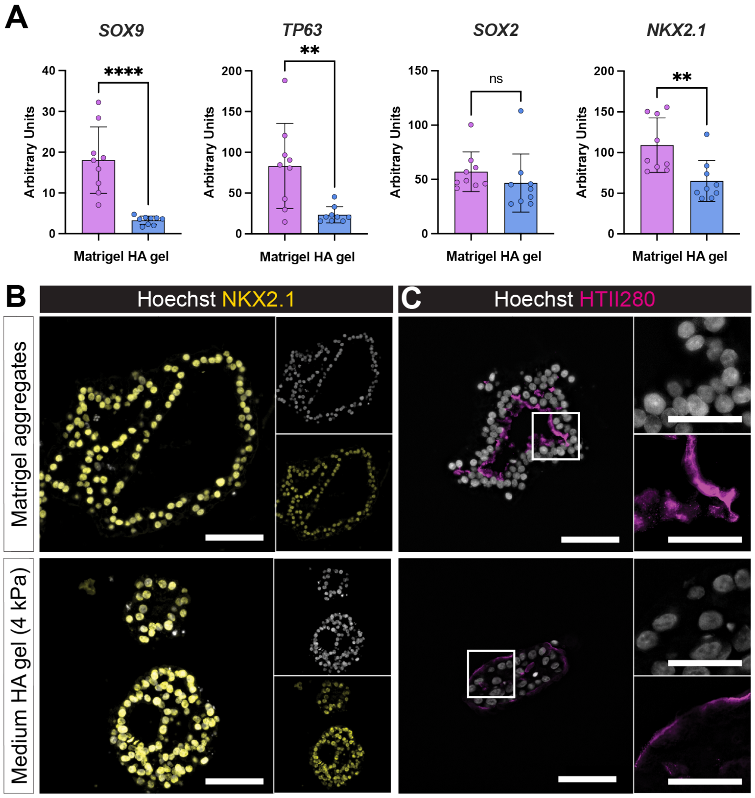


**Figure S2. Aggregates maintain AT2 identity in HA hydrogels and Matrigel**. **A**. qPCR characterization of alveolar organoids in HA hydrogels for SOX9, a marker of bud tip progenitor cells, TP63, a marker of bud tip progenitor cells, SOX2, an airway epithelial cell marker, and NKX2.1, a marker of lung epithelial identity. n = 9 repeated measurements from 3 independent experiments. **** p<0.0001, ** p<0.001, ns - no significant difference by unpaired Student's t-test. **B**. Representative immunofluorescence images of NKX2.1, a lung epithelial marker in Matrigel and HA hydrogels. Top inset, Hoechst, bottom inset, NKX2.1. Scale bar 50 μm. **C**. Representative immunofluorescence images of HTII280, an AT2 cell apical marker in Matrigel and HA hydrogels. Top inset, Hoechst, bottom inset, HTII280. Scale bar 50 μm, inset 25 μm.

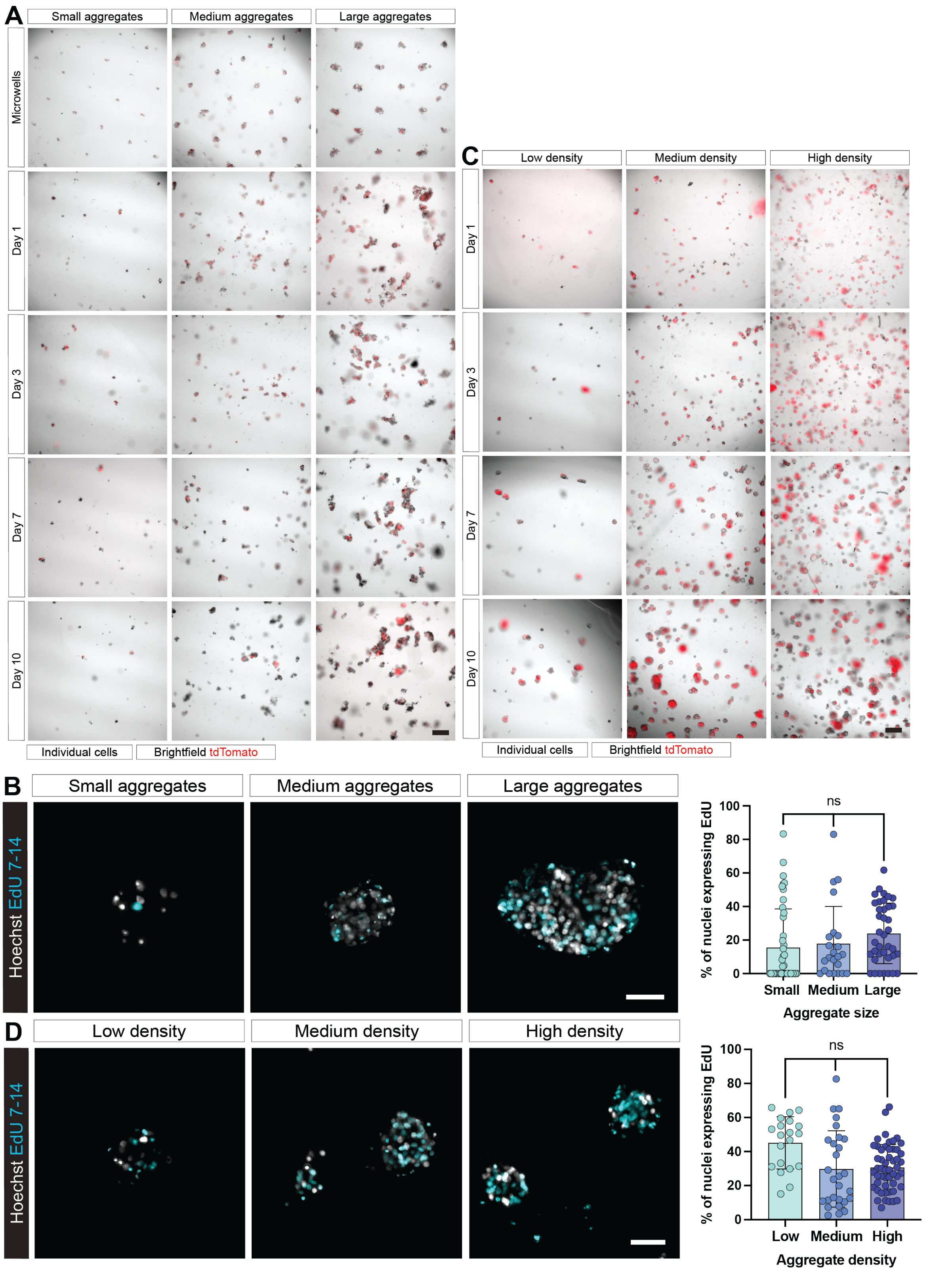


**Figure S3. Microwells enable control over aggregate size and density**. **A**. Representative images of differently sized aggregates in the microwells and in HA hydrogels for up to 10 days. Scale bar 300 µm. **B**. Representative images of EdU staining from days 7-14 of various aggregate sizes in HA hydrogels on day 14. Scale bar 50 µm. Quantification of % of nuclei expression EdU days 7-14 at various aggregate sizes. n ≥ 23 organoids from 3 independent experiments. ns - no significant difference by one-way ANOVA with Tukey's multiple comparisons test. **C**. Representative images of aggregates at different densities in HA hydrogels over the culture period. Scale bar 300 µm. **D**. Representative images of EdU staining from days 7-14 of various aggregate densities in HA hydrogels, fixed on day 14 in culture. Scale bar 50 µm. Quantification of % of nuclei expression EdU day 7-14 at various aggregate sizes. n ≥ 20 organoids from 3 independent experiments. ns - no significant difference by one-way ANOVA with Tukey's multiple comparisons test.

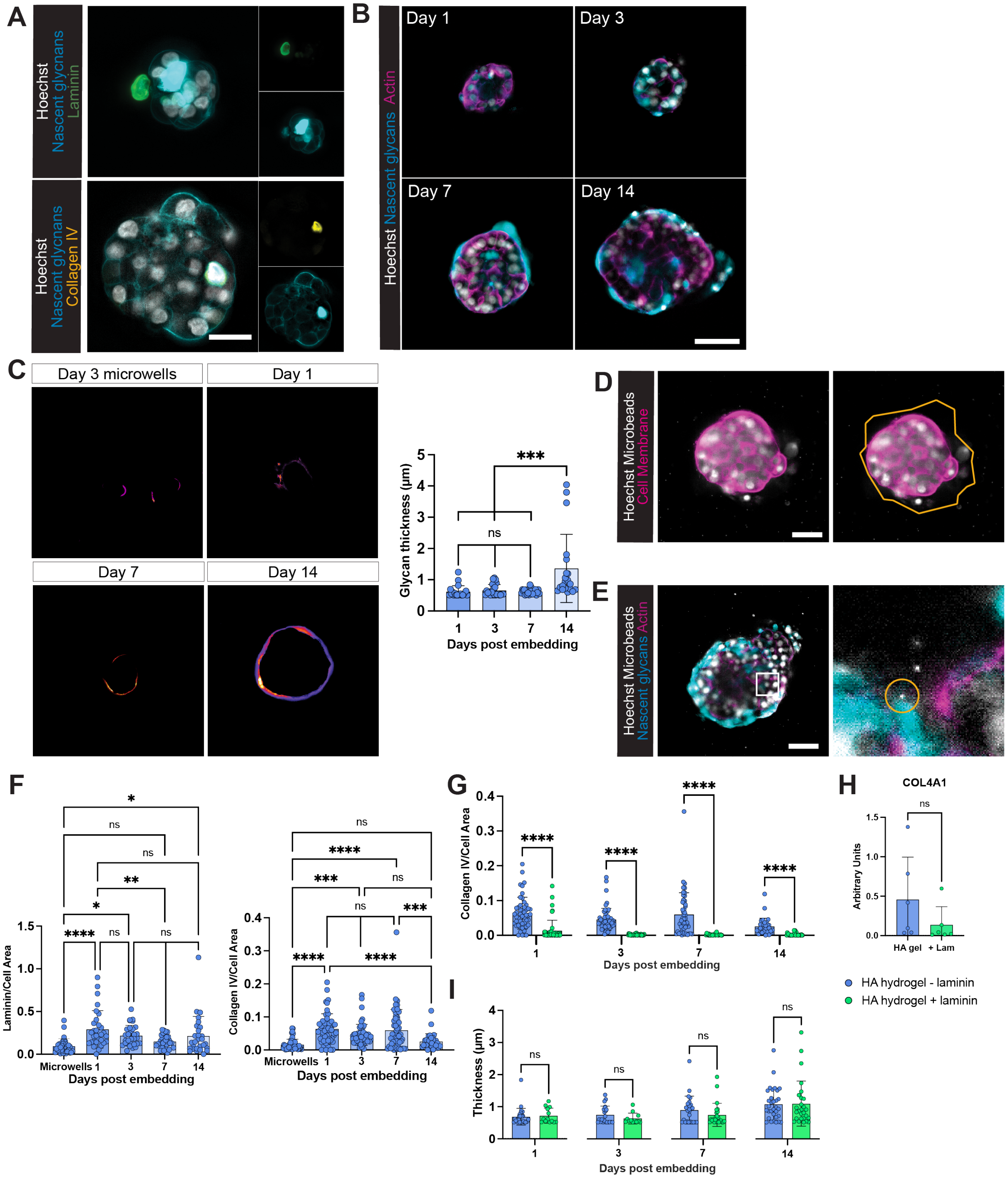


**Figure S4. Nascent ECM secretion direct aggregates and organoid function**. **A**. Representative immunofluorescence staining of basement membrane proteins laminin (green) and collagen IV (yellow) with nascent glycans (cyan) of aggregates in microwells. Scale bar 50 µm. **B.** Representative images of nascent glycans secreted by embedded aggregates on day 1 and 14. Scale bar 50 µm. **C**. Representative images and quantification of projected nascent glycan thickness generated with ImageJ BoneJ plugin. n ≥ 20 alveolar organoids from 3 independent experiments. *** p<0.001, ns - no significant difference by one-way ANOVA with

Tukey's multiple comparisons test. **D**. Schematic of hydrogel distance quantification (i.e., the area between the aggregate/organoid and the hydrogel). Beads adjacent to the organoid were traced to form a polygon, and aggregate/organoid area was subtracted from the polygon area. **E**. Representative image of day 14 organoid in HA hydrogel with fluorescent microbeads at day 14. Right, inset showing microbead at the periphery of nascent glycans (yellow circle). **F**. Quantification of laminin and collagen IV per cell area in the aggregates and throughout the culture period post-embedding. n ≥ 28 aggregates across 3 independent experiments. ns - no significant difference, * p<0.05, ** p<0.01, *** p<0.001, **** p<0.0001 by one-way ANOVA with Tukey's multiple comparisons test. **G**. Collagen IV area per cell area in HA hydrogels without laminin (blue bars) and with added exogenous laminin (green bars) across the culture period. n ≥ 28 ROI across 3 independent experiments. **** p<0.0001 by unpaired Student's *t*-test. **H**. qPCR for Col4A1 expression in HA hydrogels without laminin (blue bar) and with exogenous laminin (green) bar at day 14 in the culture period. ns - no significant difference by unpaired Student's t-test. **I**. Nascent glycan thickness in HA hydrogels without laminin (blue bars) and with added exogenous laminin (green bars) across the culture period. n ≥ 12 ROI across 2 independent experiments. ns - no significant difference by unpaired Student's t-test.

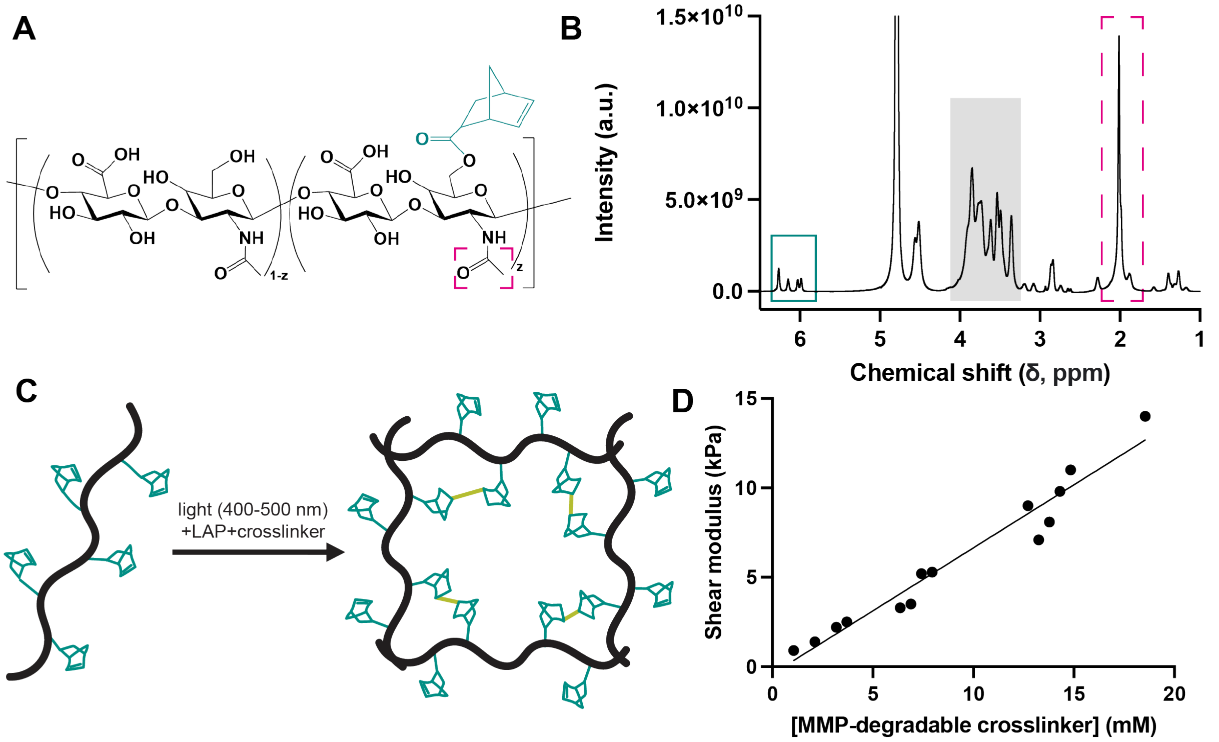


**Figure S5. HA hydrogel characterization**. **A**. Norbornene-modified HA (NorHA) polymer (chemical structure. Hyaluronic acid backbone is in black, norbornene in green. **B**. NorHA NMR. Hyaluronic acid backbone is shaded in gray, norbornene groups in solid green box, N-acetyl groups in pink dashed box. **C**. Schematic of NorHA polymer crosslinking to form a hydrogel. Thiol-ene chemistry at the norbornenes enables dithiolated sequences to act as a crosslinker and form the hydrogel. Thiolated cell-adhesive ligands can also be added to the NorHA backbone at the norbornenes. **D**. Graph of shear modulus of NorHA with varying crosslinking concentration.

### Supplementary Tables

##### Supplementary table 1. qPCR primers

| Primer name | Forward | Reverse |
|---|---|---|
| LAMP3 | GTTCTAAACGGAAGCAGACTCT | CGTTGGGGTCGATGTTGAAG |
| SFTPB | GGGTGTGTGGGACCATGT | CAGCACTTTAAAGGACGGTGT |
| SFTPA | TGTCCTCCTGGAAATGATGG | GGCTTGGAGCTCCTCATCTA |
| NAPSA | TTCCGGGGCCACACTGAT | GGTTCTCTCCATCCCCTCAG |
| SFTPC | AGCAAAGAGGTCCTGATGGA | CGATAAGAAGGCGTTTCAGG |
| NKX2.1 | CTCATGTTCATGCCGCTC | GACACCATGAGGAACAGCG |
| TP63 | CCACAGTACACGAACCTGGG | CCGTTCTGAATCTGCTGGTCC |
| SOX2 | CCATCATTGGAGCAGGAATC | GACCAGCGGTAAGATTTCCA |
| SOX9 | GTACCCGCACTTGCACAAC | GTGGTCCTTCTTGTGCTGC |
| COL4A1 | GGGATGCTGTTGAAAGGTGAA | GGTGGTCCGGTAAATCCTGG |
| RN18S | GCAGAATCCACGCCAGTACAAG | GCTTGTCCAGACCATTGGC |

##### Supplementary table 2. Antibodies

| Name | Company | Catalog # | Dilution |
|---|---|---|---|
| Primary antibodies | | | |
| Rabbit anti-SFTPC | Seven Hills Bioreagents | WRAB-76694 | 1:500 |
| Mouse anti-SFTPB | Seven Hills Bioreagents | WMAB-189 | 1:250 |
| Mouse anti-SFTPA | Leica | NCL-L-SP-A | 1:200 |
| Armenian hamster anti-MUC1 | ThermoFisher Scientific | MA50-11202 | 1:500 |
| Mouse anti-ECAD | BD Biosciences | 610181 | 1:500 |
| Rabbit anti-ZO1 | Cell Signaling Technologies | 13663 | 1:400 |
| Rabbit anti-Laminin | Abcam | 11575 | 1:500 |
| Mouse anti-Collagen IV | Invitrogen | PA5-78028 | 1:200 |
| Rabbit anti-NKX2.1 | Abcam | Ab76013 | 1:250 |
| Rabbit anti-RFP | Rockland | 600-401-379 | 1:500 |
| Mouse anti-HTII280 | Terrace Biotech | TB-27AHT2-280 | 1:100 |
| Rabbit anti-KRT5 | Sigma Aldrich | HPA059479 | 1:500 |
| Mouse anti-AcTub | Sigma Aldrich | T7451 | 1:1000 |
| Secondary antibodies | | | |
| Goat anti-mouse IgG Alexa Fluor 288 | Abcam | Ab150113 | 1:200 |
| Goat anti-mouse IgG Alexa Fluor 594 | Abcam | Ab150116 | 1:200 |
| Goat anti-mouse IgG Alexa Fluor 647 | ThermoFisher Scientific | A21235 | 1:200 |
| Goat anti-rabbit IgG Alexa Fluor 488 | ThermoFisher Scientific | A32731 | 1:200 |
| Goat anti-rabbit IgG Alexa Fluor 594 | Abcam | Ab150080 | 1:200 |

| | | | |
|---|---|---|---|
| Goat anti-rabbit IgG Alexa Fluor 647 | ThermoFisher Scientific | A21244 | 1:200 |
| AffiniPure donkey anti-mouse IgM | Jackson ImmunoResearch | 715-545140 | 1:500 |
| AffiniPure goat anti-Armenian hamster IgG Alexa Fluor 594 | Jackson ImmunoResearch | 127-585-160 | 1:500 |
